# A quantitative network modeling approach to evaluate the role of cytokine combinations on CD4+ T cell differentiation, partial polarization, and plasticity

**DOI:** 10.1101/232884

**Authors:** Mariana E. Martinez-Sanchez, Leonor Huerta, Elena R. Alvarez-Buylla, Carlos Villarreal

## Abstract

Diverse cellular polarization states with different phenotypes and functions are derived from the differentiation of activated CD4^+^ T naïve lymphocytes in the presence of particular cytokines. In addition, conversion of polarized cells to phenotypes different from that originally induced has been documented, highlighting the capacity of the immune response for adaptation to changing circumstances. In a recent study, we proposed a minimal Boolean regulatory network of CD4+ T differentiation that incorporates transcription factors, signaling pathways, and autocrine and exogenous cytokines. The qualitative model effectively reproduced the main polarized phenotypes of CD4^+^ T cells and several of the plasticity events reported in the literature. Yet, the amount and the expression of cytokines relative to expression of other factors influence CD4+ T cell transitions. In this paper, we have extended the Boolean network to a continuous model that allows us to assess the effect of quantitative differences in the concentrations and combinations of exogenous and endogenous cytokines, as well as diverse levels of transcription factors expression, in order to assess the role of intracellular and extracellular components in CD4^+^ T differentiation and plasticity. Interestingly, the model predicts either abrupt or gradual differentiation patterns between observed phenotypes depending on critical concentrations of single or multiple environmental cytokines. Plastic changes induced by environmental cytokines were observed in conditions of partial phenotype polarization in the Th1/Th2 transition. On the other hand, the Th17/iTreg transition was highly dependent on cytokine concentrations in the environment. Thus, modeling shows how the concentration of exogenous factors, the degree of initial polarization, and cell heterogeneity, may determine the differentiation and plasticity capacity of CD4^+^ T cells. The model and results presented here are useful to further understand system-level mechanisms underlying observed patterns of CD4^+^ T differentiation and plasticity.

## Introduction

The phenotype of a cell emerges from the feedback between internal regulatory networks and microenvironmental signals. The combination, concentration, and duration of the latter can either stabilize a transcriptional profile characteristic of a particular cell lineage or promote a transition into another cell type (DuPage and Bluestone, 2016; Stockinger and Brigitta, 2010). CD4+ T cells constitute a useful model to evaluate the role of micro-environmental signals and regulatory elements on cell function. In this system, the combination and concentration of exogenous cytokines are crucial for CD4+ T cell differentiation and plasticity (DuPage and Bluestone, 2016; Stockinger and Brigitta, 2010; Eizenberg-Magar et al., 2017).

CD4+ T cells are part of the adaptive immune response. Naïve CD4+ T cells are activated in response to antigens presented by antigen presenting cells (APC) (Zhu et al., 2010). Depending on the cytokines in the microenvironment, these cells may differentiate into particular subsets. APCs are the main source of cytokines (extrinsic cytokines), but they can also be produced by other cells of the organism (Duque and Descoteaux, 2014; Sozzani et al., 2017). Exogenous cytokines bind to the membrane receptors of the cell and activate intracellular signaling pathways. These signals activate or inhibit particular transcription factors and promote the production of autocrine cytokines, creating a positive feedback that reinforces the polarization dynamics (Zhu et al., 2010). In addition, autocrine cytokines can also activate or inhibit other cells of the immune system and combinations of cytokines can have synergistic or antagonistic effects on CD4+ T cell differentiation procesess that orchestrate pathogen attack, modulation of the immune response, or immunopathology (Zhu et al., 2010).

Functional CD4+ T lymphocytes can be grouped into subsets known as Th1, Th2, Th9, Th17, Treg, Tr1, and Tfh. It has been documented that Th1 cells require extrinsic IL-12 and IFNγ, they express T-bet and IFNy (Perez et al., 1995; Hsieh et al., 1993; Szabo et al., 2000, 2003). Th2 cells require extrinsic IL-4 and are stabilized by IL-2, they express GATA3, IL-4, IL-5 and-IL13 (Cote-Sierra et al., 2004; Le Gros et al., 1990; Zheng W, 1997; Ansel et al., 2006; Swain SL Weinberg AD, 1990). Th17 cells require extrinsic TGFβ and IL-6, IL-21 or IL-23, they produce RORγt, IL-21, IL-17A and IL-17F (Zhou et al., 2007; Ivanov et al., 2006; Veldhoen et al., 2006; Korn et al., 2009). Treg cells require extrinsic TGFβ and IL-2, they express Foxp3, TGFβ and in some cases IL-10 (Davidson et al., 2007; Zheng et al., 2007; Chen et al., 2003; Hori S Nomura T, 2003). Th9 cells require IL-4 and TGFβ, they express IL-9 (Schmitt et al., 2014; Kaplan, 2013; Lu et al., 2012). Tr1 and Th3 cells express IL-10 and TGFβ respectively (Roncarolo et al., 2006; Awasthi et al., 2007; Gagliani et al., 2015). Tfh cells require IL-21, they express Bcl6 (Johnston et al., 2009; Nurieva et al., 2009; Yu et al., 2009; Crotty, 2014).

Furthermore, CD4+ T cells are highly heterogeneous, suggesting that cell populations go through a continuum of polarization levels after initial priming (Stockinger and Brigitta, 2010; DuPage and Bluestone,2016; Magombedze et al., 2013; Eizenberg-Magar et al., 2017). Thus, mixed cellular phenotypes may be encountered under particular cytokine concentrations and combinations, and in some cases, hybrid cell types such as Th1-like and Th2-like regulatory cells or Th1/Th2 hybrids (Hegazy et al., 2010; Koch et al., 2009; Wohlfert et al., 2011). Studies performed on polarized CD4+ T cell populations indicate that, even under controlled in vitro conditions, stimulation generates heterogeneous cell populations with variable cytokine expression profiles or intermediate cell types (Assenmacher et al., 1994; Bucy et al., 1994; Openshaw et al., 1995; Kelso et al., 1999; Eizenberg-Magar et al., 2017). Asymmetric cell division with segregation of signaling proteins may explain this behavior (Verbist et al., 2016).

The same cytokines responsible for the induction of naïve cells to a particular polarized state may also dictate the conversion from a different subset to this state. For example, multiple studies report the transit of Treg cells towards Th17 cells in response to the addition of exogenous IL-6 in the presence of TGFβ (Yang et al., 2008; Stockinger and Brigitta, 2010; Lee et al., 2009a). Other plastic transitions depend on the degree of polarization, as in the case of the Th17/Treg (Berod et al., 2014; Michalek et al., 2011; Gagliani et al., 2015) and the Th1/Th2 transition (Perez et al., 1995; Murphy et al., 1996; Hegazy et al., 2010). Recently polarized Th1 and Th2 cells can transdifferentiate into other subsets in response to environmental IL-4 or IL-12, but fully polarized Th1 and Th2 cells are robust to changes in the microenvironment (Murphy et al., 1996). Despite abundant experimental data, understanding of the underlying mechanisms that explain the observed differentiation and plasticity patterns of these cells must require the integrated analysis of interactions between the molecular pathways induced by microenvironmental factors, and heterogeneity in the internal capacity of cells for activation.

Complex regulatory networks give rise to stable multidimensional configurations (attractors) that correspond to expression profiles that, in this case, characterize specific CD4+ T cell subsets (Kauffman, 1969; Mendoza et al., 1999; Bornholdt, 2008; Villarreal et al., 2012; Martínez-Sosa et al., 2013; Albert and Thakar, 2014; Naldi et al., 2015; Álvarez-Buylla et al., 2016). Multidimensional Epigenetic Landscapes (EL), originally proposed as a metaphor, can be studied as a dynamical system determined by a set of coupled non-linear interactions between the network nodes that result in the emergence of a diversity of steady-state patterns; each expression pattern may be attained in a number of ways starting from different initial configurations of the expression levels of the network components. This number determines the size of the attraction basin, and consequently, the probability that a given pattern is expressed is proportional to this size. (Cortes et al., 2008; Villarreal et al., 2012; Davila-Velderrain et al., 2015). Even though the mathematical structure of the network interactions does not allow the derivation of an explicit potential function representing the EL, a probabilistic approach to the EL (not considered in this paper) is still possible (see Villarreal, 2012). On the other hand, the analyses presented here reveals the conditions required to drive the system from one attractor to another one (Haken, 1977). Indeed, the steady states (or attractors) of the dynamical system, may be visualized in as states lying at the bottom of valleys of the EL. From this point of view, we explore not only pathways that conduce to equilibrium valleys, but also alterations of the expression levels of components of the network that induce the transit between adjacent valleys.

By considering the former approach, we explore how the restrictions imposed by the regulatory network determine or affect the patterns of CD4+ T cell differentiation and plasticity (Mendoza, 2006; Naldi et al., 2010; Carbo et al., 2013; Abou-Jaoudé et al., 2014; Martinez-Sanchez et al., 2015; Eizenberg-Magar et al., 2017). With that purpose, we propose a continuous version of an already considered regulatory model (Martinez-Sanchez et al., 2015). The continuous model enables evaluations of quantitative alterations of the inputs (exogenous cytokines) and the intrinsic components (transcription factors, signaling pathways, autocrine cytokines) of the network by using a model of ordinary differential equations (Villarreal et al., 2012; Davila-Velderrain et al., 2015). The study contemplates a method specifically designed to study the EL repatterning under altered microenvironmental conditions (Davila-Velderrain et al.,2015; Pérez-Ruiz et al., 2015). In general, this type of systemic dynamic multi-stable models constitutes useful tools to integrate experimental data and may provide novel predictions concerning phenotypic patterns in Systems Biology. Our simulation results reproduce the main polarized phenotypes of CD4+ T cells and several of the plasticity patterns reported in the experimental literature. We determine the effect of systematic changes in the concentrations of exogenous cytokines and the internal state of the network in the differentiation and plasticity of CD4+ T cells. We focus on the Th1/Th2, and Th17/iTreg transitions, which have been thoroughly characterized given their therapeutic relevance. In addition, our study provides some novel interesting predictions that may be tested experimentally.

## Methods

Similarly as the original Boolean network, its continuous variant includes nodes that correspond to transcription factors, signal transduction pathway components, cytokine receptors, SOCS proteins, as well as autocrine and exogenous cytokines [File S1]. The edges of the network correspond to the verified regulatory interactions between the nodes [Figure 1, File S2] (Martinez-Sanchez et al., 2015). In contrast with the Boolean approach, the expression level of each node may acquire continuous values within the range (0,1) and the network interactions are characterized by fuzzy logic propositions. In fuzzy logic (Novak et al.,1999), the degree to which an object exhibits a property is given by a membership function (specified below) so that the truth values of variables may range between completely false (0) and completely true (1). These extreme values correspond in our system to the basal expression level (inactive) and to the maximum expression level (active) of a node, respectively. Fuzzy propositions may be built by replacing ordinary logical operations by fuzzy connectors, so that logical conjunction *A AND B* corresponds to the ordinary product *A*\**B*, logical disjunction *A OR B* to *A*+*B*-*A*\**B*, and logical negation *NOT A* to *1-A*. This methodology, discussed in Ref. (Azpeitia et al., 2014) has been used successfully for several other systems (See review in Davila-Velderrain et al., 2017, and allows to construct a parameter-free model that evidences the regulatory logics of the system and yields the observed biological behavior. The model was validated by verifying that the predicted CD4^+^ T subsets and plasticity transitions coincide with experimental observations [Figure 1, File S3].

**Figure 1.**
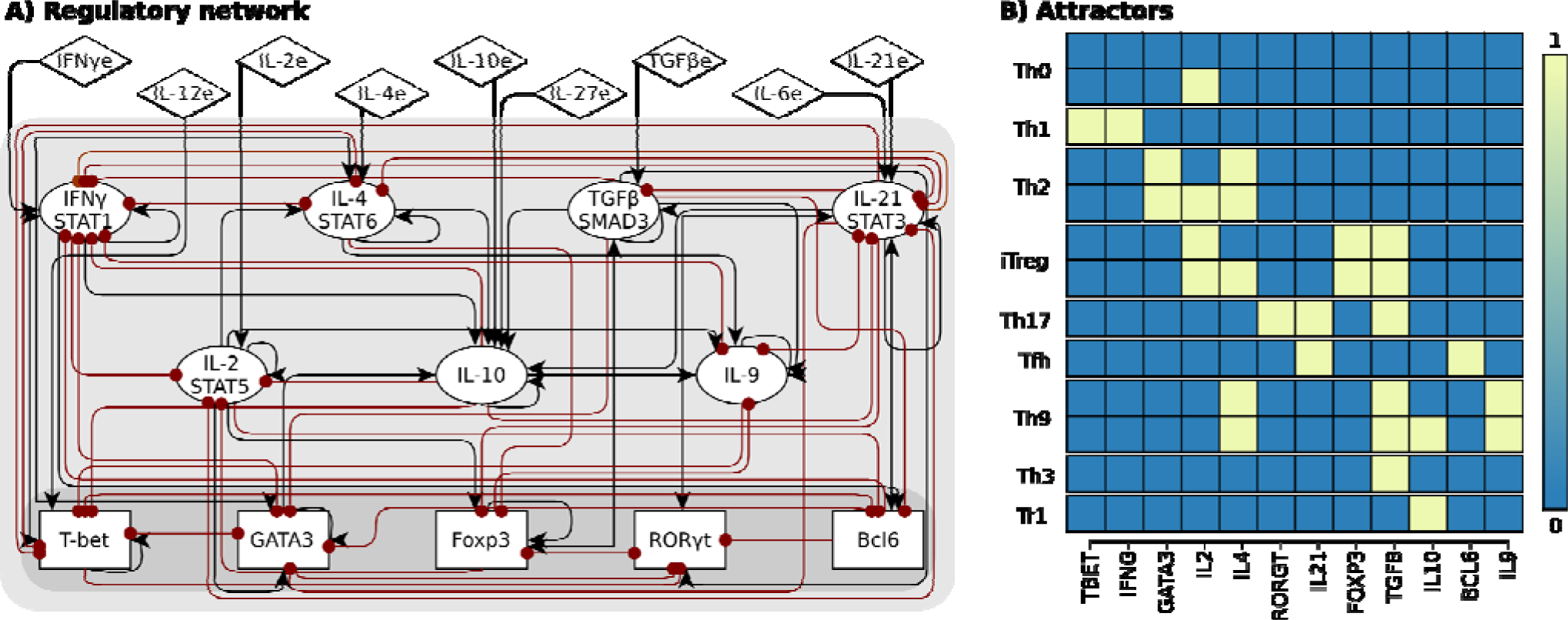
CD4+ T cell transcriptional-signaling regulatory network. The regulatory network was constructed using available experimental data. (A)The network includes transcription factors (rectangles), autocrine cytokines and their signaling pathways (ellipses) and exogenous cytokines (diamonds). Interactions leading to activation are represented by black arrows, while those leading to inhibition with red dots. (B) Sample attractors of the system.

The simulations performed in this work consider a reduced version of the extended T CD4+ regulatory network presented in file S1 Fig. of Ref. (Martinez-Sanchez et al., 2015). Reduction was achieved by either collapsing downstream linear signaling pathways of the regulatory network, or by using logical algebraic rules (Naldi et al., 2009; Villarreal et al., 2012). The final network consisted of 21 nodes [Fig. 1). Five nodes correspond to transcription factors (TBET, GATA3, FOXP3, RORGT, and BCL6); seven nodes correspond to signaling pathways integrating signal transducers such as STAT proteins, interleukin receptors, and autocrine cytokines (IFNg, IL2, IL4, IL10, TGFB, IL9, and IL21); nine nodes correspond to exogenous cytokines, that are produced by other cells of the immune system and that serve as inputs to the network (IFNGe, IL12e, IL2e, IL4e, IL10e, IL27e, TGFBe, IL6e and IL21e). These are marked with an “e” after the cytokine name. To study the effect of the microenvironment we focused on nine biologically relevant environments (Zhu et al., 2010): pro-Th0, pro-Th1, pro-Th2, pro-Th17, pro-Th9, pro-Tfh, pro-iTreg, pro-Tr1, and pro-Th3 [Table 1]. The regulatory cytokine IL-10 deserves special consideration, since it uses STAT3, similarly as the inflammatory cytokines IL-6 and IL-2. Thus, we assume that IL-10 signaling is mediated by an independent pathway than IL-6/IL-21, even though they share STAT3 as a messenger molecule (Moore et al., 2001). While IL-27 has been linked to multiple functions, we consider that its main role in the model is a regulatory cytokine (Pot et al., 2009; Murugaiyanet al., 2009; Awasthi et al., 2007). The model ignores weak interactions, chemokines, and epigenetic regulation that are also relevant and should be included in future modeling efforts.

**Table 1:** **Exogenous cytokines in different environments included in the CD4+ T cell regulatory network.** Active nodes refer to the same exogenous cytokines, whose concentrations were modified during the simulation, adopting values between 0 and 1.

| Environment | Active input nodes |
| --- | --- |
| pro-Th0 | None |
| pro-Th1 | IFNγe, IL12e |
| pro-Th2 | IL2e, IL4e |
| pro-Th17 | IL21e, TGFBe |
| proTh9 | IL4e, TGFBe |
| proTfh, | IL21e |
| pro-iTreg | IL2e, TGFBe |
| pro-Tr1 | IL10e, IL27e |
| pro-Th3 | TGFBe |

If we now assume that *q_i_*(*t*) represents the expression level of node *i* at a given time, its rate of change determined by the regulatory network interactions may be described by

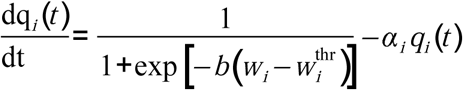

Here, the first term on the right hand side represents a (logistic) membership function discussed above. It depends on the fuzzy logic proposition w_i_, with a threshold value w_i_^thr^ that renders the proposition as true. The parameter b is a rate indicating the rapidity of the transition from false to true. Finally,α_i_ is a decay rate, so that if the membership degree is null, the node expression level decays exponentially at this rate. In this paper we suppose that α_i_=1, w_i_^thr^=1/2, and b=25. The resulting attractors of the dynamical system are presented in [File S4].

The continuous model enables evaluations of quantitative alterations of the inputs (exogenous cytokines) and the intrinsic components (transcription factors, signaling pathways, autocrine cytokines) of the network. To investigate the polarization process we assume an initial state that corresponds to a CD4^+^ T cell activated under non-polarizing cytokine conditions (Th0) where we suppose that nodes representing transcription factors, signaling pathways or autocrine cytokines are initially inactive. In order to model plastic transitions, we consider a cell in an initial polarization state determined by different expression levels of the characteristic transcription factor and cytokines [File S3] in the range 0≥q_i^int^_≥1. In both kinds of experiments, we represent the effect of the microenvironment using a selected set of exogenous cytokines [Table 1] that are active with relative concentrations lying in the range 0≥q_i^ext^_≥1. For each initial condition and relative concentration of the microenvironment, we simulate the system to obtain the final steady state and measure the Euclidean distance between the initial state and the final steady state.

We consider that a steady state of the system corresponds to a CD4^+^ T cell subset if its characteristic transcription factor and cytokines [File S5] are actively expressed. Given the continuous range of expression levels in the system, we introduce a criterium to define node expression. We consider that a node is actively expressed if q_i_≥ 0.75, unexpressed if q^i^ ≤0.25, while intermediate values correspond to a transition zone, with no definite expression. We consider that an attractor corresponds to a given subset if the characteristic cytokines and transcription factors are active. On these terms, a cell was classified as Th0 if it expressed low levels (q_i_< 0.25) of transcription factors [File S5].

We observe that the polarization transitions between steady states of the system display two alternative behaviors as a function of exogenous cytokine concentrations. In the first case, a sudden (discontinuous) transition appears at a critical concentration of polarization inducing cytokines. In the second case, the transition is gradual (continuous), involving a transition region with intermediate expression levels of endogenous components, which could be interpreted in terms of phenotype heterogeneity. The code for all the simulation experiments performed in this work is available in [File S6].

## Results

### Network construction

A previous study reported a data-based Boolean regulatory network for CD4+ T cell differentiation and plasticity [File S1 and S2]. While the Boolean model correctly reproduces CD4+ T cell fate attainment, it is unable to determine how the different expression levels of the components of the network (transcription factors, signaling pathways, autocrine and exogenous cytokines) affect CD4+ T cell differentiation and plasticity. To address this, we converted the equations to a continuous dynamical model [File S4] (Mendoza,2006; Villarreal et al., 2012), and used the EL reshaping methodology (Davila-Velderrain et al., 2015) to address which combinations and quantitative alterations of cytokine and transcription factors expression levels are sufficient for CD4+ T differentiation and plasticity.

The stable states or attractors to which the regulatory network converges are interpreted as the characteristic expression profiles of particular CD4+ T cell subsets or types (Mendoza et al., 1999;Naldi et al., 2015; Albert and Thakar, 2014; Bornholdt, 2008; Azpeitia et al., 2011; Álvarez-Buylla et al., 2016). Given the continuous nature of the regulatory network model presented here, it is impossible to determine all the possible steady states. It was verified that the steady states of the discrete model could be recovered in the continuous model, and they were classified into CD4+ T cell subsets. It was considered that a steady state corresponded to a subset if it expressed a high concentration (qi> 0.75) of the characteristic cytokine and transcription factor associated with the subset, which in this work corresponded to Th0, Th1, Th2, Th17, Treg, Tfh, Th9, Tr1, and Th3. We also recovered steady states that correspond to GATA3+IL4-, Th1-like and Th2-like regulatory cells, which had been previously reported (Martinez-Sanchez et al.,2015; DuPage and Bluestone, 2016). Steady states that had intermediate values were considered to be in a transition zone (t.z.) of phenotypic coexistence.

### Effect of the concentration of the extrinsic cytokines in the differentiation of CD4^+^ T cells

To evaluate how the concentrations of the exogenous cytokines in the environment shapes CD4^+^ T cell differentiation, we studied the activation process of a Th0 cell as a function of increasing concentrations of the exogenous cytokines and determined the final steady states [Figure 2]. The exogenous cytokines IL12e, IFNGe, IL4e, IL6e, IL21e, TGFBe, and IL10e induce the differentiation from a Th0 initial steady state towards Th1, Th2, Tfh, Th3, and Tr1 subsets respectively (Zhu et al., 2010). On the other hand, Th17, Th9, and iTreg subsets were not induced by a single exogenous cytokine in the micro-environment. These observations are consistent with the requirement of TGFBe in combination with IL6e/IL21e, IL4e or IL2e to induce these subsets. (Zhu et al., 2010).

**Figure 2.**
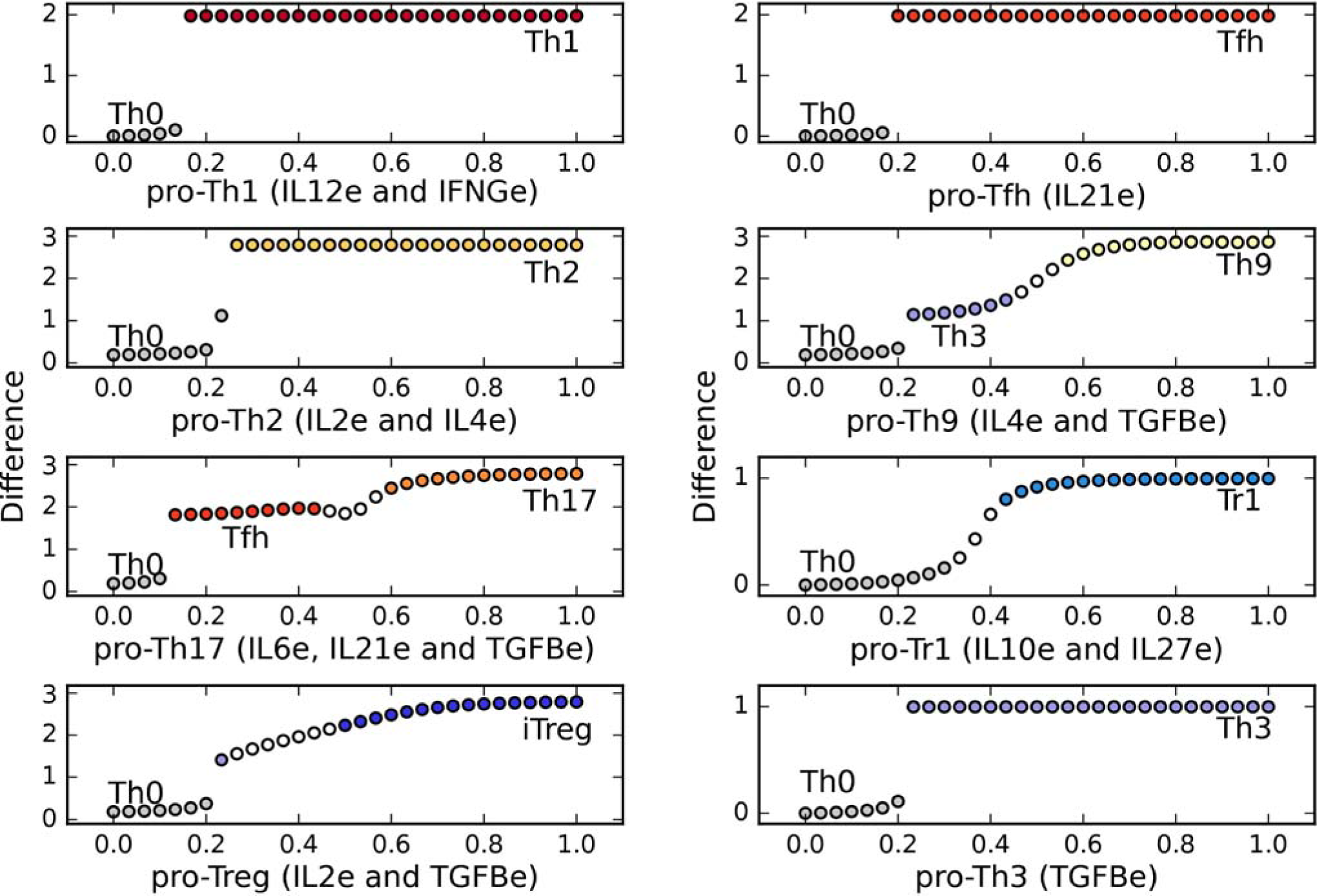
CD4+ T cell fate as a function of the concentration of single exogenous cytokines: IL12, IFNG, IL2, IL4, IL6, Il21, TGFB, IL10, IL27. From an initial state TH0, a CD4+ T cell may acquire diverse phenotypes on an abrupt or gradual transition, depending on critical concentrations of environmental cytokines. We observe that the presence of either IL12 or IFNg is sufficient for Th1 polarization, as well as IL4, is sufficient for TH2 polarization. On the other hand, IL2 alone does not lead to an effector phenotype. Similarly, the presence of either IL6 or IL21 alone is sufficient for Tfh induction, as is the case of TGFB and IL10, leading to Th3 and Tr1, respectively. IL27 alone does not lead to any fate transition in this model.

The critical concentration required to induce a transition varied depending on the particular exogenous cytokine. IL12e, IL6e, and IL21e required relatively small concentrations (0.2) to induce the differentiation from Th0 to Th1 and Tfh respectively, while IL4e required a higher concentration (0.36) to induce the differentiation from Th0 to Th2. On the other hand, IL2e and IL27e alone were not able to induce transitions. We observed that IL2e induced the expression of high levels of IL2; however, we labeled the resulting cells as Th0, as IL-2 production by itself is not associated with a particular polarization subset.

It is also interesting to note that transitions among subsets have different patterns of sensitivity to exogenous cytokine concentrations. Most of the transitions from Th0 to other subsets were discontinuous; once a threshold concentration was achieved, the cell changed its expression pattern to a different one in an abrupt manner. An exception was observed when IL10 was used as an inducer. This cytokine caused a gradual transition from Th0 to Tr1; in this case, a continuous range of steady states is attained in the transition zone between both subsets. These results show that, for most of single cytokines, CD4^+^ T cells would initiate differentiation once the threshold concentration has been reached, whereas they may display a range of sensivities to the concentration of other cytokines to induce differentiation into alternative phenotypes.

CD4^+^ T subsets such as Th9, Th17, and iTreg require combinations of cytokines to differentiate from naïve cells. In our model, we simulated the activation of a Th0 cell in the presence of different combinations and concentrations of the exogenous cytokines associated with the microenvironment [Table 1, Figure 3]. In the case of requiring more than one exogenous cytokine, all the implicated nodes were set to the same value. Using this methodology, we were able to induce the differentiation from a Th0 steady state towards Th1, Th2, Th17, Th9, Tfh, iTreg, Th3, and Tr1 subsets by cytokine combinations that are in agreement with experimental data (Zhu et al., 2010, Crotty2014, DuPage2016c).

**Figure 3.**
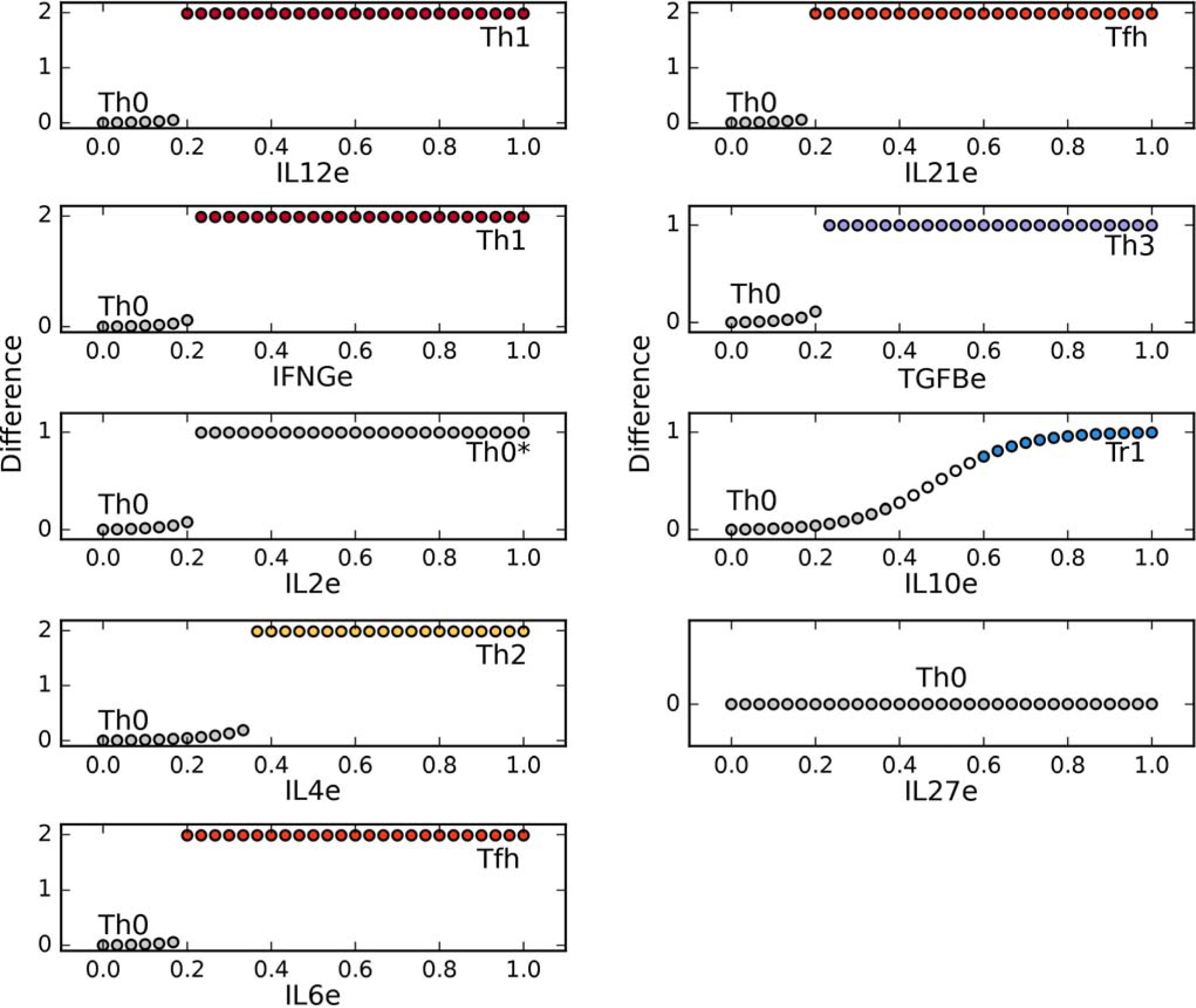
T-CD4 cell fate as a function of exogenous cytokine concentrations defining diverse phenotype-associated environments. From the TH0 initial state, a T CD4 cell evolves to different phenotypes, depending on critical concentrations of environmental cytokines as shown in Table 1: Th1 (IFNG and IL12), Th2 (IL4, Il2), Th17 (Il21, TGFB), Treg (IL2, TGFB), Tfh (IL21), Th9 (IL4, TGFB), Tr1 (IL10, IL27), Th3 (TGFB). The transition may be abrupt or gradual and, interestingly, may involve an intermediate state, as in the cases Th0 -> Tfh -> Th17 (C), and Th0 -> Th3 - > Th9 (F).

The concentration required to induce transitions when using multiple cytokines varied depending on the CD4^+^ T cell type, but it was always lower [Figure 2] than if only a single exogenous cytokine was added. This result suggests that the underlying CD4^+^ T cell differentiation regulatory network mediates a synergistic effect of cytokines on CD4^+^ T cell differentiation. For example, while a concentration of IL4e = 0.36 was necessary to induce the polarization towards Th2, a concentration of IL 2e and IL4e = 0.26 was sufficient to induce the same transition. Similarly, while a concentration of IL10e = 0.6 was necessary to induce the polarization towards Tr1, a concentration of IL10e and IL27e = 0.43 produced the same transition. Furthermore, autocrine IL10 achieved its maximum value with a lower concentration of exogenous cytokines when IL10e and IL27e act synergistically.

The transitions in pro-Th1, pro-Th2, and pro-Tfh microenvironments were abrupt, while the transition in a pro-Tr1 environment was gradual. In a pro-Th17, pro-Th9, and pro-iTreg micro-environments, all including TGFβe, there was a small abrupt change followed by a gradual change in the expression values of the steady state. Also, in pro-Th17 and pro-Th9 we found an intermediate step before the final polarized state. In the pro-Th17 case, increasing cytokine levels induced an initial abrupt change towards a plateau zone corresponding to Tfh, followed by a transition to the Th17 steady state. A similar behavior was observed in the pro-Th9 microenvironment with a precursor TGFβ^+^ (Th3) subset, followed by a final Th9 steady state. It is worth noting that TGFβ has a key role in the induction of the three types of CD4^+^ T cell types discussed here trough complex relationship with the effect of other exogenous cytokines (Eizenberg-Magar et al., 2017). These results illustrates the capability of the minimal model to contribute to the understanding of how the context determine the cellular response to TGFβ in the immune system.

Thus, the continuous model describes the cytokine concentration dependence of the differentiation of T cell subsets, illustrating abrupt or gradual changes, the effect of cytokine combinations and, notably, the induction of different subsets under the action of different concentrations of the same cytokine combinations.

### Effects of the concentration of the exogenous and endogenous cytokines and transcription factors in on the plasticity of CD4^+^ T cells

Once differentiated, CD4^+^ T cells may undergo plastic transitions to other subsets under specific conditions. As discussed above, in order to model this kind of transitions we studied phenotypic alterations of cells stated in a defined polarization state, as a function of different concentration levels of exogenous cytokines able to induce an alternative phenotype.

We first focus on the transition between Th1 and Th2 [Figure 4], which has been experimentally observed, particularly when these cells have recently differentiated (Perez et al., 1995; Panzer et al., 2012). To study this process we considered the response of already differentiated Th1 and Th2 states, to the presence of variable concentrations of the of the subset-defining cytokine in combination with the opposing cytokine (IFNGe for Th2, and IL4e, for Th1), and determined the final steady state. When the initial configuration of the system corresponded to a highly polarized Th1 (TBET and IFNG = 1) or Th2 (GATA3 and IL4 = 1) states, for every combination of (exogenous) IL4e and IFNGe concentrations, the system remained in its original state even under high concentrations of these cytokines, respectively, indicating that highly polarized Th1 or Th2 cells are not plastic. However, by considering initial lower concentrations of Th1 and Th2 transcription factors and cytokines, consistent with partial phenotype polarization, plastic transitions ensue. CD4^+^T cells require the production of high levels of autocrine IFNG and expression of TBET to maintain a Th1 phenotype. If the expression levels decrease, especially in the case of autocrine IFNG; it will transit into a Th2 cell. At the same time, the cells require the production of high levels of autocrine IL4 and expression of GATA3 to maintain a Th2 phenotype. If the initial expression levels decrease it will transit into a Th1 cell. At high expression levels of initial GATA3 and low initial IL4, there exists a transition zone where the cell displays mixed characteristics. These results show that plasticity between the Th1 and Th2 subsets depends not only on the cytokines present on in the microenvironment but also on the intracellular state.

**Figure 4.**
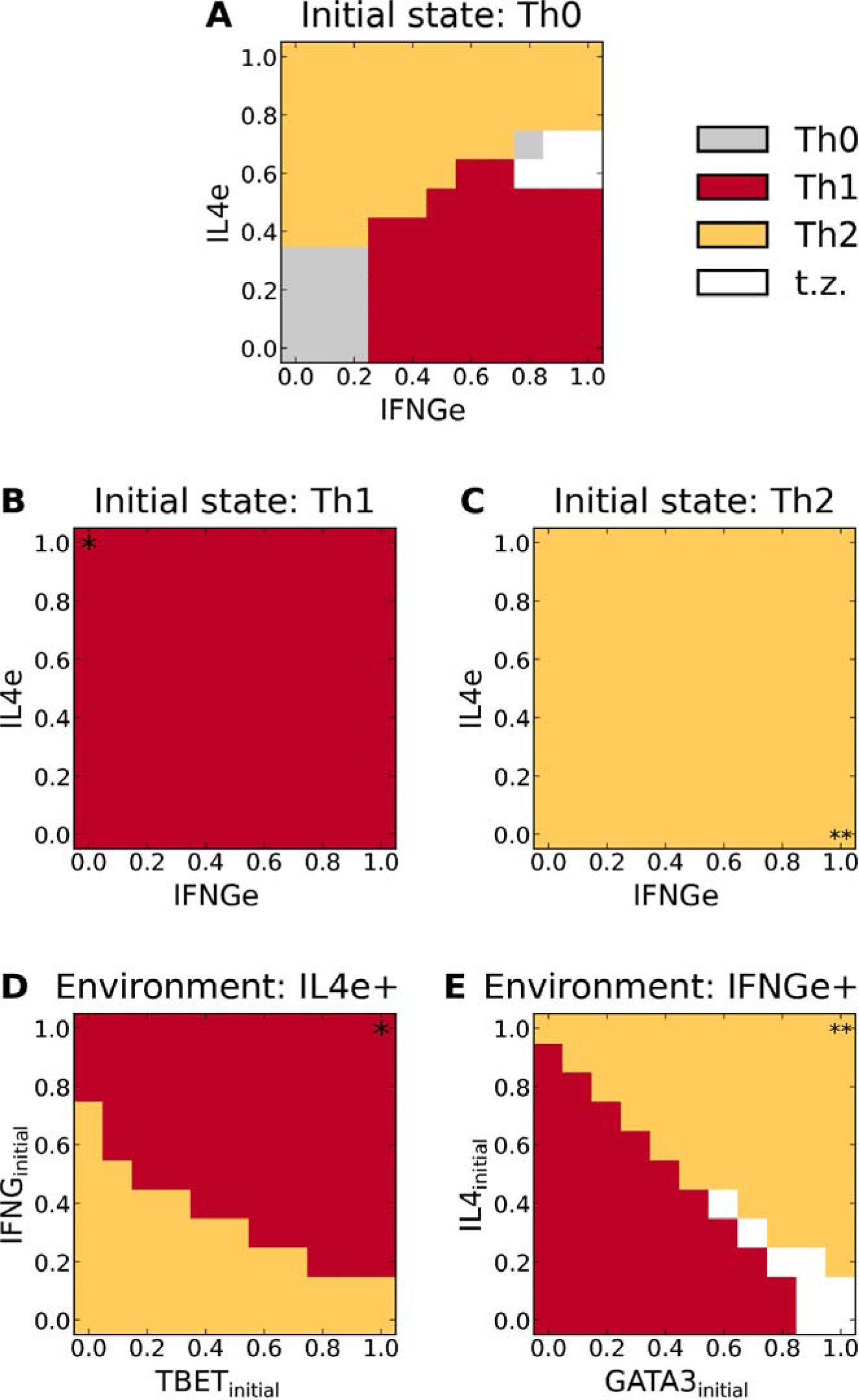
Phenotype space diagrams for Th1 and Th2 polarization and plasticity as a function of the relative concentration of environmental IFNg and IL4, and expression of transcription factors. (A) Diagram for cell differentiation assuming an initial Th0 state. As the external concentration of IFNGe increases, the system develops an abrupt transition from Th0 to Th1. Similarly, an increase in external IL4e drives an abrupt transition from TH0 to Th2. For moderate concentrations of IFNGe and IL4e (< 0.8), we observe two wide zones of Th1 or Th2 prevalence with a sharp boundary, meaning that small variation of cytokines at this zones may change cell polarization. A transition zone with no defined polarization appears at higher concentrations of these cytokines (white and gray areas). (B and C) Plasticity diagrams assuming full Th1 (B) or Th2 (C) polarized states (i.e., induced by INFg= 1 and IL-4= 1 in diagram A, respectively). No phenotypic transitions are observed under variable concentrations of environmental IL4e and autocrine IFNGe. (D) Plasticity diagram of Th1 cells assuming an environmental concentration of IL4e = 1. Cells require the production of initial high levels of autocrine IFNG and expression of TBET to maintain a Th1 phenotype. If the initial expression levels decrease, especially in the case of autocrine IFNG, it will transit into a Th2 cell. (E) Plasticity diagram of Th2 cells assuming an environmental concentration of IFNGe = 1. The cell requires the production of high levels of autocrine IL4 and expression of GATA3 to maintain a Th2 phenotype. If the initial expression levels decrease it will transit into a Th1 cell. At high expression levels of initial GATA3 and low initial IL4, there exists a transition zone where the cell cannot be classified.

The transition between Th17 and iTreg, has been extensively investigated experimentally (Lee et al., 2009b; Littman and Rudensky, 2010; Wei et al., 2008; Xu et al., 2007; Noack and Miossec, 2014; Leeet al., 2009a; Kleinewietfeld and Hafler, 2013) [Figure 5] since this plasticity case is particularly important for some pathological conditions, such as chronic inflammation. To study this process we considered fully differentiated Th17 (RORGT and IL21=1) and iTreg (FOXP3 and TGFB = 1) under the presence of different concentrations of the exogenous cytokines IL2e, IL21e, and TGFBe. In the case of Th17 cells, they remained in a Th17 phenotype at a high concentration of TGFBe, while they switched towards Tfh for lower concentrations of TGFBe (< 0.6). Some experiments have reported that induction of Th17 require exogenous TGFB (Veldhoen et al., 2006), but it is uncertain if the transition towards Tfh associated to low TGFB levels will occur in all cases. On the other hand, iTreg cells remain stable provided that high concentrations of IL2e are present, while they transit towards Th17, Tfh or Th3 when concentrations of IL2e are low (< 0.65). This shows that the plastic transitions between subsets are not symmetrical, and depend on the previous polarization state of the cell.

**Figure 5.**
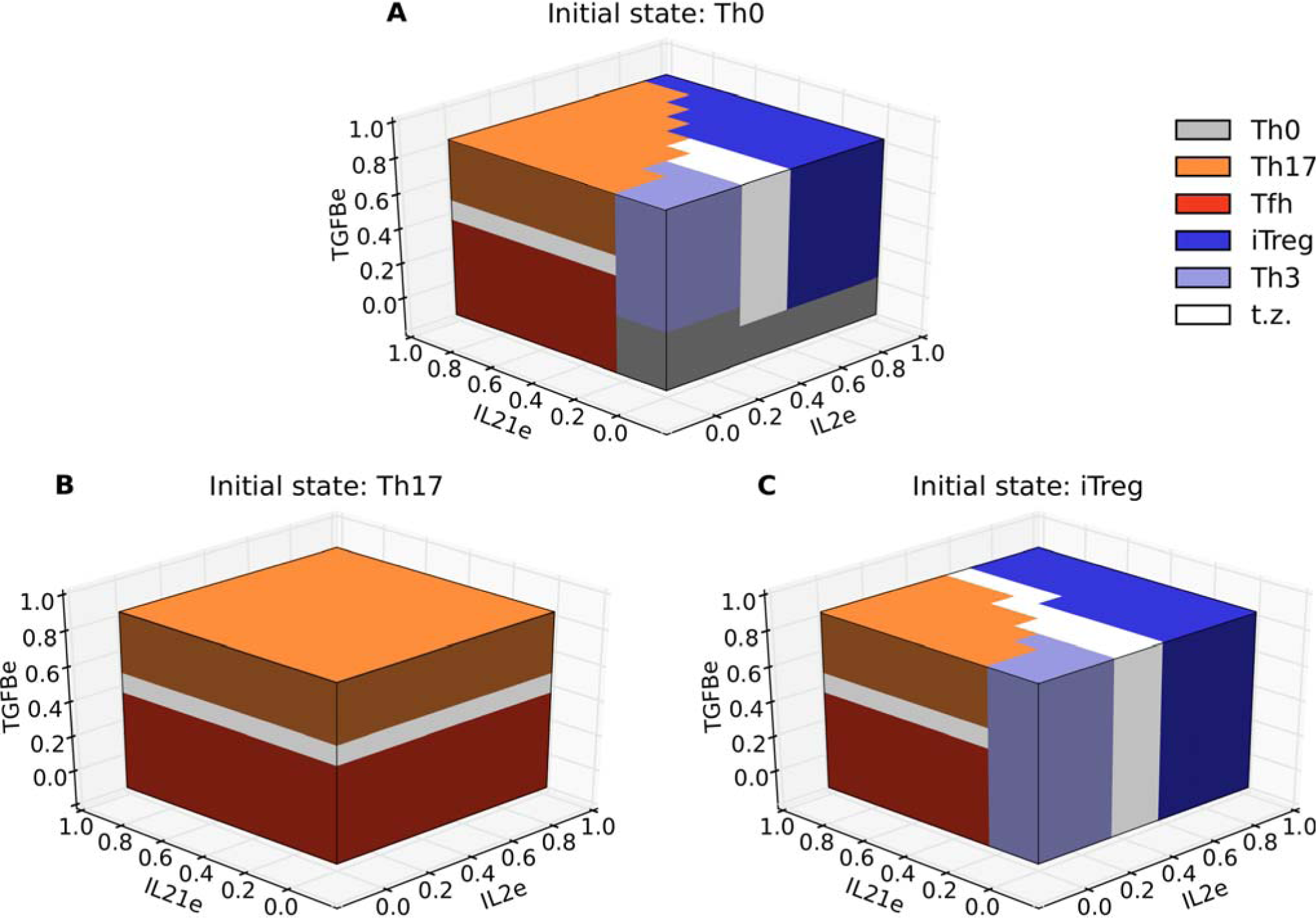
Three-dimensional phenotype space diagrams for Th17 and iTreg polarization and plasticity as a function of the relative concentrations of IL2, IL21, and TGFB in the microenvironment. In the differentiation diagram (A) we observe alternative phenotypic regions defined by relative concentrations of environmental cytokines. The regions may be either separated by a sharp boundary or by a more gradual transition zone (labeled in white). The plasticity diagram (B) indicates a polarized behavior for Th17 versus Tfh phenotype determined by a high or low concentration of external TGFB. A richer behavior ensues when the initial state is Treg, as shown in the plasticity diagram (C). We observe a similar structure as that depicted in (A), except that the Th0 zone is absent.

## Discussion

Our simulations show different profiles of differentiation patterns of CD4+ T cells that depend on the concentration and combinations of exogenous cytokines, highlighting the importance of synergy and competing interactions at the signaling network level. We also showed that plasticity between the Th1/Th2 and iTreg/Th17 subsets can be obtained by varying the concentration of microenvironmental cytokines and the expression level of transcription factors and autocrine cytokines derived from the internal state of the cell.

The model predicts both abrupt and gradual transitions between cell types. In abrupt transitions, there is a sudden change in the final steady state or cell type, once the concentration of exogenous cytokines exceeds a threshold value. This behavior suggests that the transition between stable cell phenotypes is energetically favorable once the threshold value has been achieved. In this process, exogenous cytokines provide the initial stimulus to promote the expression of both transcription factors and autocrine cytokines in positive feedback loops that increase the stability of attractors to quickly attain polarization.

In contrast, in gradual transitions, steady states that express intermediate levels of transcription factors and autocrine cytokines appear. In these steady states, a clear-cut threshold between the two expression patterns is not observed, so they cannot be easily classified into one subset or another, signaling the manifestation of partially polarized states. The heterogeneity of CD4+ T cells has been well-documented (DuPage and Bluestone, 2016; Stockingerand Brigitta, 2010; Eizenberg-Magar et al., 2017), and could be the result of regulatory circuits capable of generating a range of cells that express intermediate levels of specific molecules and that can stably coexist or change from one another under certain conditions. It is important to notice that every gradual transition involves regulatory circuits with central nodes which display feedback interactions. This feedback renders stability to the initial polarization state so that its intrinsic cytokine production and transcription factor expression should gradually decrease under changing microenvironmental conditions. We observed this behavior especially in response to changes in the concentration of IL-10 and TGFβ. IL-10 is a regulatory cytokine produced by multiple CD4+ T subsets (Gagliani et al., 2015; Howes et al., 2014). TGFβ may display both regulatory and inflammatory effects and it is implied in the differentiation of multiple subsets like Th17, iTreg, and Th9 (Veldhoen et al., 2006; Davidson et al., 2007; Chen et al., 2003; Kaplan, 2013). It is conceivable that gradual transitions and generation of intermediate polarization states reflect the intricate regulatory signaling effects of TGFβ of IL-21, which may participate in a biological mechanism of tuning their effect in the immune response (Grossman & Paul, 2015).

The model also captures some cases where there is an abrupt transition followed by a gradual transition in polarization processes. Such is the case of the Th0-Tfh-Th17, the Th0-Th3(TGFB+)-Th9 and the Th0-iTreg transitions. Interestingly, in all these cases TGFβ is present in the micro-environment. This indicates that the concentration of TGFβ may modulate the immune response in complex ways, putting forward a system-level explanation of experimental results. It is known that TGFβ regulates Th17 cells in a differential way depending on the concentration and cytokines in the environment (Yang et al., 2008). Furthermore, consistent with our simulations, it is known that the TGFβ signaling pathway is highly modulated (Attisano and Wrana, 2002; Travis and Sheppard, 2014). Our model also predicts that that TGFβ may induce distinct subsets at different concentrations, in particular, Tfh, Th9, iTreg, and Th3. Studying this effect would require polarization experiments with careful control of extracellular TGFβ concentration.

The model highlights the cooperation between exogenous cytokines during differentiation. Th17, iTreg, and Th9 subsets require TGFβ in combination with IL-6/IL-21, IL-2 and IL-4 to differentiate, respectively, in agreement with experimental data (Veldhoen et al., 2006; Davidson et al., 2007; Chen et al., 2003; Kaplan, 2013). In other cases, the effect of a single cytokine is sufficient to induce polarization, but the synergy with other cytokines lowers the threshold concentration necessary to induce polarization. In this way, the model allows us to quantitatively study and predict synergistic relations among cytokines in CD4+ T cell differentiation.

As mentioned above, we also use the model to study the effect of opposing cytokines in the differentiation and plasticity of Th1/Th2 and Th17/iTreg subsets. The Th1 and Th2 cells are highly stable, and the transition between them is hard to achieve experimentally (Perez et al., 1995; Murphy et al., 1996; Hegazy et al., 2010). We found that, once they have achieved a stable state, Th1 and Th2 are robust to changes in their microenvironment. However, partially polarized cells can transit to the other cell types when they are subject to an opposing cytokine (IL-4 in the case of Th1 or IFNγ in the case of Th2). These results agree with the observation that recently polarized Th1 and Th2 cells are plastic, but fully polarized are not (Murphy et al., 1996). This behavior seems consistent with a particularly robust interaction circuit, defined by four regulatory switching modules between mutually inhibitory nodes with negative feedback, each node defining an alternative regulatory route.

On the other hand, iTreg and Th17 cells are less stable, and there are multiple reports of single cell transitions caused by changes in the environment, especially from iTreg to Th17 cells (Lee et al.,2009b; Littman and Rudensky, 2010; Wei et al., 2008; Xu et al., 2007; Noack and Miossec, 2014; Lee et al., 2009a; Kleinewietfeld and Hafler, 2013). Our model shows that the Th17 phenotype is stable in the presence of TGFβ, whereas it transits towards Tfh cells in the absence of exogenous TGFβ. While this transition has been reported (Crotty, 2014), we cannot test its stability in the long term, as the model lacks many of the niche signals that stabilize Tfh cells \cite{Crotty2014, DuPage2016c}. Modeling also shows that maintenance of the iTreg phenotype requires high concentrations of IL-2, which is in accordance with experimental observations (Veldhoen et al., 2006; Davidson et al., 2007; Chen et al., 2003), although they transit towards Th17 in the presence of high concentrations of IL-21 and low concentrations of IL-2. Thus, the model provides a simulation of the spontaneous transition of iTreg into Th17 in the presence of IL-21 or the closely similar IL-6 (here considered as equivalents) (Xu et al., 2007) at low concentrations of IL-2. The plasticity of this transition is not symmetrical, as changes in the microenvironment are not enough for Th17 to transit towards iTreg. For such transition, it is also necessary to alter the internal state of the cell, changing the expression levels of key transcription factors, as has been shown in experimental studies (Berod et al., 2014; Michalek et al., 2011; Gagliani et al., 2015). These results seem to imply that the basin of attraction of iTreg is shallower than that of Th17. This could be the result of the different regulatory circuits implied in the differentiation of each cell type, since while both depend on TGFβ, iTreg both requires and inhibits the production of IL-2 (Pandiyan et al., 2007; Fontenot JD Gavin MA, 2003), restricting the stability of this cells.

The model and simulations presented here are able to describe cell type transitions and do not rely upon specific parameter estimates. However, the exact transition points may change depending on the precise concentrations and parameters of the biological system (Eizenberg-Magar et al., 2017). In our model, the value 0 corresponds to a basal level of expression, not to the absolute absence of the transcription factor or cytokine. Given the relative nature of the quantitative variations introduced in the model, we should be cautious in providing precise quantitative predictions concerning the sensitivity of the different subsets under real experimental conditions.

The model and simulations presented here predict two main kinds of transitions between attractors: abrupt transitions, characterized by well-defined transcriptional profiles, and gradual transitions, with a zone associated to heterogeneous populations. We propose that the second kind is associated with feedback regulatory interactions as well as switching modules involved in the dynamics of the system. A careful analysis of this kind of regulatory circuits will shed light on the specific mechanisms defining transcriptional programs that lead to cell heterogeneity. This is especially important in the case of TGFβ, which has a crucial role in the regulatory network. Understanding the interactions underlying the dynamical behavior may help elucidate the regulatory role of this important molecule in the immune response.

Theoretical models like the one presented here provide an ideal tool to integrate recent advances in experimental knowledge and provide a system-level mechanistic explanation for observed behaviors in experiments, and also to provide informed predictions for future experiments. Hence, the feedback between experimental and theoretical research is necessary to understand the rich behavior of CD4+ T cells and the immunological system.

## Author Contributions

ERAB and CV conceived, planned and coordinated the study. CV and MEMS established the continuous model and performed simulations and calculations. All of the authors participated in the interpretation, result analysis, and wrote the paper.

## Acknowledgments

We acknowledge Diana Romos for her support with logistical tasks. We thank Jose Davila-Velderrain and Juan Arias del Angel for providing code for this project.

